# Transmitter Co-Expression Reveals Key Organizational Principles of Local Interneuron Heterogeneity in the Olfactory System

**DOI:** 10.1101/167403

**Authors:** Kristyn M. Lizbinski, Gary F. Marsat, Andrew M. Dacks

## Abstract

Heterogeneity of individual neurons within a population expands the computational power of the entire neural network. However, the organizing principles that support heterogeneity within a neuronal class are often poorly understood. Here, we focus on a highly heterogeneous population of local interneurons whose traits co-vary seemingly at random. We asked if local interneurons (LNs) in the antennal lobe (AL) of *Manduca sexta* express fixed, predictable combinations of neurotransmitters, or if transmitter co-expression can be explained by random probability. We systematically determined the co-expression of neuropeptides and GABA by LNs and found variable patterns of co-expression for all neuropeptides, except for tachykininergic LNs which exhibited highly stereotyped co-expression on a neuron-by-neuron basis. To test if observed patterns of co-expression were random, we used a computational model and found that the probabilities of transmitter co-expression cannot be explained by independent expression of each transmitter. We also determined that setting a single rule in the model, while leaving the rest of the co-expression up to random probability, allowed the model to replicate the overall heterogeneity of transmitter co-expression across antennal lobe LNs. This implies that certain co-expression relationships contribute to the ground plan of the AL, but that otherwise, transmitter expression amongst LNs may be random, allowing heterogeneous co-expression patterns to emerge. Furthermore, neuropeptide receptor expression suggests that peptidergic signaling from LNs may simultaneously target olfactory receptor neurons, LNs and projection neurons, and thus the effects of different peptides do not segregate based on principal AL cell type. Our data suggest that while specific constraints may partially shape transmitter co-expression in LNs, a large amount of flexibility on a neuron-by-neuron basis produces heterogeneous network parameters.

## Introduction

Neurons are typically categorized into classes by obvious morphological similarities, physiological properties, or relative location. However, each neuronal class can be surprisingly heterogeneous in their synaptic, biophysical and transcriptional profiles (Cohen et al., 2015; Eddine et al., 2015; Okaty et al., 2015). Heterogeneity of individual neurons within a population expands the computational power of the entire network. For example, local interneurons (LNs) as a class tend to be particularly heterogeneous, such as cortical (Flames and Marin, 2005) and hippocampal interneurons (Maccaferri and Lacaille, 2003) and the insect antennal lobe (AL) is no exception (Nassel and Homberg, 2006; Chou et al., 2010; Seki et al., 2010). Much of the computational capacity of the AL is mediated by LNs that refine the information transferred between olfactory receptor neurons (ORNs) and output neurons (projection neurons; PNs). LNs perform many tasks including presynaptic gain control, divisive normalization, and dynamic control of response range (reviewed in (Martin et al., 2011) and (Wilson, 2013)). Furthermore, LNs are highly diverse and vary in their morphology, physiology (Chou et al., 2010; Seki et al., 2010; Reisenman et al., 2011), and transmitter co-expression (Homberg et al., 1990; Berg et al., 2007; Utz et al., 2008; Carlsson et al., 2010; Siju et al., 2013; Fusca et al., 2015).

Since LNs often express more than one transmitter, the presence of specific sets of neuropeptides or combinations of transmitter types allows neurons to have a combinatorial impact on their targets (Nusbaum et al., 2001; Tritsch et al., 2016; Nusbaum et al., 2017). However, neuropeptides are often only partially co-expressed across the entire LN population (Utz et al., 2008; Fusca et al., 2015), opening the possibility that the expression of individual transmitters may be determined independent of each other or even randomly. In the crustacean stomatogastric ganglion, varying synaptic strength, membrane properties and conductance at the level of a single neuron still produces similar and robust network output (Prinz et al., 2004; Marder, 2011). Thus network dynamics permit a considerable amount of variability in individual physiological parameters, as long as reliable output can be produced from animal to animal. Furthermore, there are several examples of biological systems in which specific features, like gene expression (Elowitz et al., 2002; Raj and van Oudenaarden, 2008; Huh and Paulsson, 2011) or anatomical layout (Caron et al., 2013), appear to be randomly structured. For example, random combinations of AL PNs from different glomeruli converge and synapse upon individual mushroom body Kenyon cells in *D. melanogaster* regardless of anatomy, developmental origin or odor tuning, thus abandoning the odor-topic organization of the AL (Caron et al., 2013). Similarly, it could be that co-expression in LNs is highly variable, or seemingly random, and that specific, tightly bound co-expression patterns are not necessary. However, the ground rules for patterns of LN transmitter co-expression are unknown.

Here, we took a systematic approach to understanding the heterogeneity of transmitter co-expression within the olfactory system of *M. sexta*. LNs in the AL of *M. sexta* co-express a wide array of transmitters like GABA and neuropeptides (Homberg et al., 1990; Utz et al., 2007; Utz et al., 2008). We envision two extreme scenarios to explain the observed co-expression patterns. Each transmitter could be expressed independently of each other in a subset of cells and co-expression levels would reflect the probability that a neuron express both transmitters by chance. At the other extreme, the fact that a neuron expresses transmitter A determines the probability it also expresses transmitter B (i.e. their expression is not independent). To determine where the AL lies along this spectrum, we paired immunohistochemistry with computational modeling to ask whether observed transmitter co-expression patterns could be replicated using random probability, and if not, how many co-expression rules must be specified to explain the overall heterogeneity of LNs. Finally, we identified the expression of 5 putative neuropeptide receptors and the GABA_B_ receptor across all principal neuron types of the AL to determine the network targets of peptidergic signaling via LNs.

## Results

The antennal lobe (AL) of *M. sexta* contains stereotyped cell clusters that house PNs and LNs. The lateral cell cluster (LCC) consists of ~950 cell bodies, including 590 PNs and all 360 total LNs (Homberg et al., 1988), of which 164 are GABAergic (Hoskins et al., 1986). A previous study in LNs of *M. sexta* showed that there are no relationships between physiological properties, morphological properties, or GABA expression patterns in LNs (Reisenman et al., 2011). We therefore took a systematic approach to determine if transmitter co-expression patterns could explain the apparent heterogeneity of LN cellular properties. We first determined the pairwise co-expression relationships for GABA and five neuropeptides, TKK, FMRFamide (FMRF), Allatotropin (ATR), Myoinhibitory peptide (MIP) and Allatostatin (AST) in LNs (Fig. 1A-E). Peptidergic LNs predominantly co-expressed GABA (Fig. 1F, Table 1), suggesting that LNs can be broadly subdivided into GABAergic/peptidergic and non-GABAergic/non-peptidergic LNs. The non-GABAergic LNs are likely glutamatergic as RT-qPCR on LCC mRNA revealed that the vesicular glutamate transporter (vGLUT) was highly expressed relative to a reference gene (40s ribosomal protein s3; RpS3). A large population of glutamatergic LNs in *M. sexta* would be consistent with the organization of the *D. melanogaster* AL (Das et al., 2011; Liu and Wilson, 2013). We then determined the co-expression patterns of every pairwise combination of TKK, FMRF, ATR and MIP (Fig. 2A-G). Most LNs co-express multiple neuropeptides to a variable degree (Fig. 2 E, G, C). The exception to this rule was TKK which co-localized 100% with MIP and never co-localized with FMRF and ATR (Fig. 2A, B, D, H). Co-expression ratios for each pairwise co-expression relationship (i.e. ATR with MIP) revealed that apart from TKK, neuropeptides were co-expressed to a variable degree (Fig. 2H).

**Figure 1:**
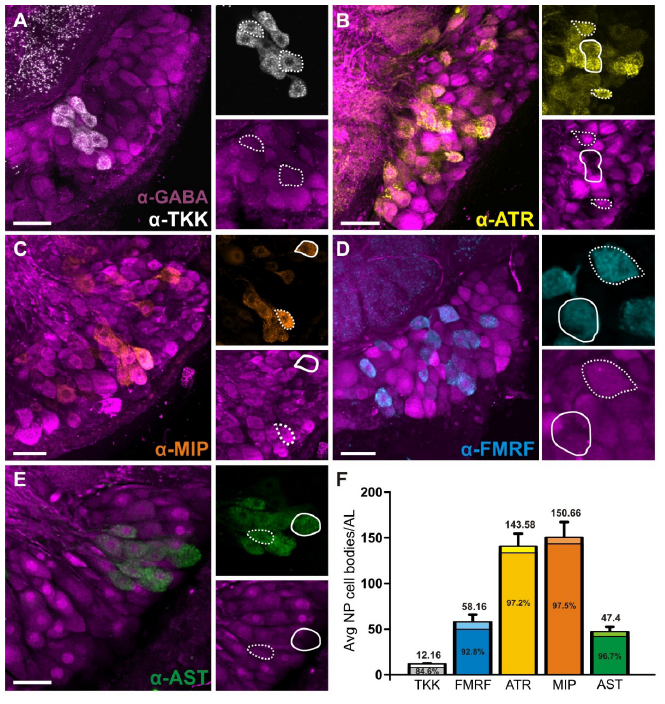
GABA and neuropeptide co-expression in the LCC. Dashed lines = co-expressed. Solid lines = not co-expressed. **A:** LCC labeled for GABA (magenta) and TKK (white). **B:** LCC labeled for GABA (magenta) and ATR (yellow). **C:** LCC labeled for GABA (magenta) and MIP (orange). **D:** LCC labeled for GABA (magenta) and FMRFamide (cyan). **E:** LCC labeled for GAB A (magenta) and AST (green). **F:** Bar graph of total number of cell bodies (above bars) that express each transmitter type per AL and the percentage (within bars) of each neuropeptide population per AL that co-express GABA. See Table 2 for averages and standard deviations. n = 6 animals per transmitter. All scale bars 50um.

**Figure 2:**
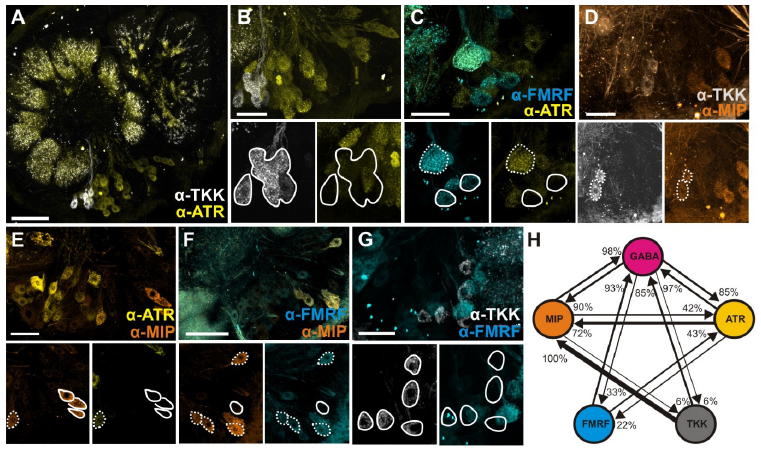
Neuropeptides are co-expressed to varying degrees. Co-expression for **A,B:** TKK(white) and ATR (yellow), **C:** FMRFamide (cyan) and ATR, **D:** TKK and MIP (orange), **E:** ATR and MIP, **F:** FMRFamide and MIP and **G:** TKK and FMRFamide. Dashed lines= co-expressed. Solid lines= not co-expressed. All scale bars = 50um.H: Schematic representation of transmitter co-expression by LNs. Each circle represents the population of LNs that express a given transmitter. Arrow width and percentage located at arrowhead represent proportion of a given LN type (arrow origin) that also express a second transmitter (arrow destination). FMRFamide and MIP co-expression could not be calculated for technical reasons (see methods). No TKK LNs co-expressed FMRFamide or ATR. Non-GABAergic LNs are not depicted.

**Table 1:** Neuropeptide cell body totals and % co-expression with GABA

| Neuropeptide | Avg # of cell bodies in LCC | % co-expression with GABA |
| --- | --- | --- |
| TKK | 12.16 ± 0.55 | 84.6% |
| FMRF | 58.16 ± 17.48 | 92.8% |
| ATR | 143.58± 24.38 | 97.2% |
| MIP | 150.66 ± 16.79 | 97.5% |
| AST | 47.4 ± 12.83 | 96.7% |

Two possible scenarios can explain the lack of apparent systematic association between specific neuropeptides co-expressed by LNs. The first possibility is that expression of a given neuropeptide is independent of the expression of another and the likelihood of specific co-expression patterns can be calculated probabilistically as the result of random co-expression. The second alternative explanation is that specific pairs of neuropeptides are co-expressed more (or less) often than by chance, and a certain number of such fixed relationships can explain the overall pattern of neuropeptide co-expression. LNs are highly diverse in their physiology, transmitter profile and connectivity (Chou et al., 2010; Reisenman et al., 2011), and at first glance, neuropeptide co-expression in *M. sexta* LNs appears variable, if not random on a neuron-by-neuron basis. We therefore used computational modeling to determine if transmitter co-expression was truly random or if fixed relationships are required to explain the observed likelihood of co-expression (Supp. Fig. 1A). Given the known total number of LNs in the LCC, and the total number of LNs expressing each neuropeptide, the model calculated the random probability of a given neuron co-expressing two transmitters (see methods for full description of model). We then compared our observed ratios of transmitter co-expression to the predicted patterns from the model that assumed random probability of co-expression for each pairwise relationship (Supp. Fig. 1B). If co-expression ratios from our ‘random probability’ model matched the observed co-expression ratios, we would assume that transmitter co-expression in the LCC can be explained probabilistically. We used a standard deviation index (SDI) to measure how well our model replicates our known biological co-expression patterns (see methods for detailed description). For instance, an SDI score of 0 would indicate that our simulation perfectly recapitulates our observed biological co-expression patterns, whereas an SDI score above 1 would indicate poor performance of our model. We found that random probability alone could not explain the pattern of co-expression, as most co-expression relationships were not replicated and the model had a weighted SDI of 1.49 (Fig. 3A).

**Figure 3:**
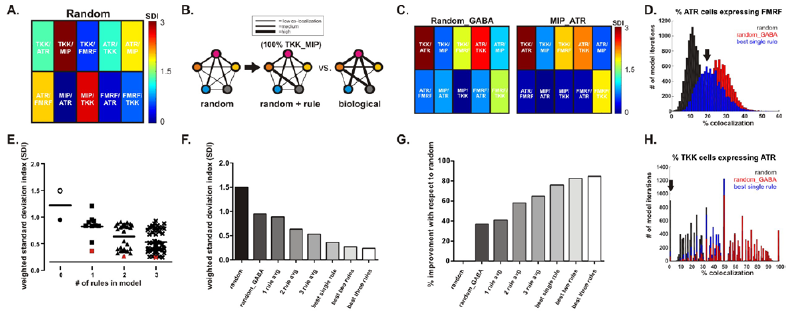
Computational analysis of transmitter co-expression reveals that random probability alone cannot explain LN heterogeneity. **A:** Random probability cannot fully rcplicate LN co-expression patterns. Each square represents an individual pair-wise relationship (e.g TKK/ATR). An SDI of 0 (blue) denotes no difference between biology and simulation, whereas values above 1 indicate a poor match between the model and biological values. **B:** The second iteration of the model followed known rules of co-expression to determine if certain relationships between transmitters could shift an otherwise randomly determined population to rcplicate known biological co-expression patterns. **C:** When the total number of neurons in the model were reduced to the total number of GABAergic LNs (a relative predictor of peptide co-expression), the model replicated more relationships than the random probability. The best single predictor of co-expression across all relationships was the rule between MIP/ATR. TKK rules were never replicated probabilistically **D:** Neither random, nor random_GABA models reliably replicated co-expression patterns. However, the MIP/ATR constraint best replicates biological co-expression patterns (denoted by black arrow). **E:** Specific rules outperform others at replicating co-expression patterns. Open circle represents model run with total # of LNs set to 360. Closed symbols represent models runs with total # of neurons set to the total # of GABAergic LNs (180). Red denotes standout iterations of the model that best replicated co-expression. **F:** Model improves as more rules are added, but MIP/ATR rule stands out. Best single rule: MIP/ATR. Best two rules: MIP/ATR + TKK/ATR. Best three rules: TKK/ATR + ATR/MIP + FMRF/ATR. **G:** % improvement of each model with respect to the random model. Both GABA constraint and MIP/ATR rule drastically improved the model’s ability to replicate co-expression patterns **H:** TKK co-expression could only be replicated by models including rules for TKK co-expression relationships.

Based on weighted SDI values calculated from the random model and our in vivo biological findings, our results indicate that neuropeptide co-expression patterns cannot be explained by combining the probabilities of independently expressed transmitters. Thus, we asked whether introducing specific co-expression rules (e.g. 100% of TKK neurons express MIP) could cause an otherwise random model to reliably replicate all co-expression relationships in the network (Fig. 3B). We first constrained the total number of cells in the model to the total number of GABAergic LNs, as presence of GABA is a reliable predictor of peptide expression. This constraint outperformed the random model and had a weighted SDI of 0.94 (Fig. 3C). This suggests that much of the diversity of neuropeptide co-expression can be constrained to the sub-population of GABAergic LNs in our study. However, of the 10 possible, single rules, applying the constraint that 42% of MIPergic LNs must co-express ATR yielded the lowest weighted SDI (0.36), far outperforming any single constraint and was well above the 95% confidence interval for all single rules (Fig 3C). Neither purely random, nor random_GABA models reliably replicated all co-expression relationships. However, the MIP/ATR constraint best replicates all co-expression patterns except those including TKK (Fig. 3C). This model also allowed for a large amount of co-expression to still be left to random chance, as only the MIP/ATR co-expression rule was specified. This suggests that the ground plan of the AL may be partially shaped by specific co-expression patterns (i.e. MIP/ATR), but that some of the heterogeneity of co-expression may be random.

No constraint produced a model that replicated TKK co-expression accurately based on random probability alone (Fig. 3H), which is likely due to TKK LNs following a strict all-or-none neuropeptide co-expression pattern. This suggests that TKK expression may be independent of the other co-expression patterns and developmentally, the co-expression patterns of TKK LNs may be more tightly controlled than other LNs or perhaps under different transcriptional regulation. TKK LNs as a population are highly stereotyped, as 100% of TKK LNs co-express MIP and 84% co-express GABA. Although there are only 12 TKK LNs (Lizbinski et al., 2016), their unique co-expression pattern suggests a distinct role from other LNs in the network.

To determine if specific combinations of rules outperformed other combinations, we provided the constrained model with every combination of 2 and 3 neuropeptide co-expression rules (for a total of 94 different model iterations). As expected, the ability of the model to replicate known biological co-expression patterns improved as more rules were added, as shown by weighted SDIs from all model iterations (Fig. 3E). However, as with constraints based on single co-expression rules, some combinations outperformed others. For example, models that included rules about MIP/ATR and TKK/ATR co-expression, were the best replicators of overall co-expression (Fig. 3E). Replicating biological co-expression patterns did not require all co-expression relationships to be fixed. This suggests that the AL may only need a few, key organizational principles that govern the observed patterns of co-expression, leaving the rest of the co-expression patterns up to random chance. Overall, both the GABA and the MIP/ATR constraint drastically improved model performance and, unexpectedly, the MIP/ATR constraint alone outperformed most 2- and 3-rule models (red square Fig. 3E and Fig. 3F-G). All other rules made incremental changes to the replicative power of the model, supporting the idea that the diversity of peptide expression examined in this study can be constrained to GABAergic LNs, and that once a single co-expression relationship is taken into account, the heterogeneity of LN neuropeptide co-expression can be explained by a random distribution.

It could be that tightly regulated co-expression of neuropeptides in specific LNs is unnecessary simply because specific classes of AL neurons express different sets of neuropeptide receptors. Thus, the impact of individual peptides within a modulatory cocktail of many peptides would be segregated due to neuron class-specific expression of each receptor. This would indicate that heterogeneous co-expression by individual LNs may not matter functionally, as long as the network relies on cell class specific receptor expression. For instance, if ORNs express the MIP receptor and PNs express the ATR receptor, an LN that co-expresses MIP and ATR could differentially target ORNs and PNs. We, therefore, asked whether neuropeptide receptors for the peptides examined in this paper were differentially expressed across the principal neuron types of the AL. We identified transcripts from the *M. sexta* genome with high sequence identity to neuropeptide receptors identified from reference genomes in closely related species (Table 2). Using RT-qPCR we determined the relative expression of 5 neuropeptide receptors (ATR, MIP, AST, FMRF, TKK), and the GABA_B_ receptor in the antennae (which house ORNs), the medial cell cluster (which houses PNs), the LCC (which houses LNs and PNs) and whole brains (as a positive control). Although the receptors for ATR, MIP, AST, FMRF, GABA_B_, were detected in all 4 tissue types, the TKK receptor was not detected in the LCC (Fig. 4, for raw qPCR data see Table 3). This suggests that TKKergic LNs differ from other LNs both in their co-expression patterns and their postsynaptic targets.

**Figure 4:**
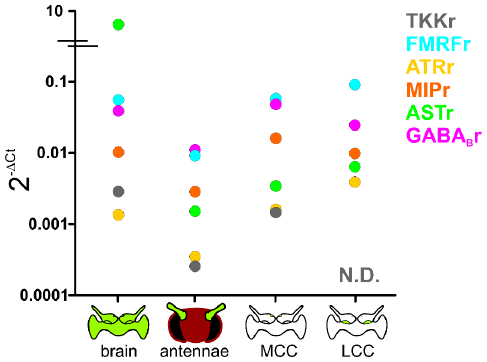
Neuropeptide and GABA_B_ receptor expression dictate functional consequences of transmitter co-expression. **A:** Relative receptor expression for ATRr, MIPr, ASTr, FMRFr, GABA_B_r are present in all tissue types and therefore expressed in ORNs, LNs and PNs in varying expression levels. TKK was not detectable (N.D.) in lateral cell cluster mRN A and therefore not detectable in LNs.

**Table 2:**
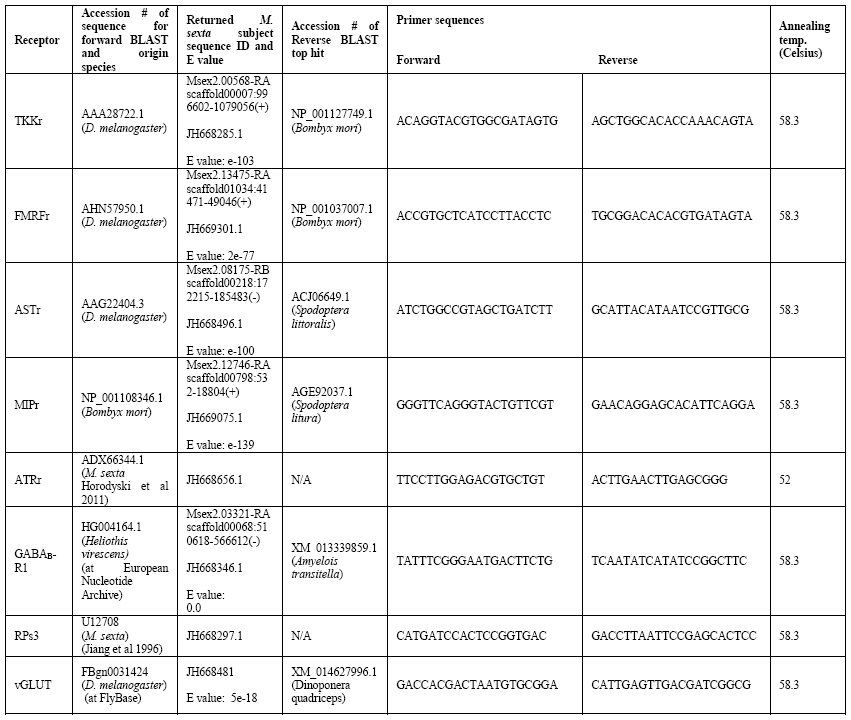
BLAST results for neuropeptide receptor primer design and primer sequences

## Discussion

Heterogeneity of individual neurons within a population expands the computational power of the entire neural network. The link between heterogeneous response properties and neural coding has been studied in a wide range of systems (Chelaru and Dragoi, 2008; Marsat and Maler, 2010; Ogawa et al., 2011; Pitkow and Meister, 2012; Ahn et al., 2014), but the role of heterogeneous neuromodulatory influence of diverse neurons in a population is not well understood. Here, we focus on a highly heterogeneous population of LNs whose traits co-vary seemingly at random. We systematically illustrate that transmitter co-expression in LNs cannot be explained by independent expression of each transmitter. However, setting a single rule in the model (MIP/ATR co-expression) while leaving the rest of the co-expression up to random chance, allowed the model to replicate the overall heterogeneity of transmitter co-expression across antennal lobe LNs. Furthermore, we found that a single neuropeptide has the potential to simultaneously target ORNs, LNs and PNs based on the expression of 4 putative neuropeptide receptors across all principal neuron types of the AL, and thus the effects of different peptides do not segregate based on principal AL cell type. Our data suggest that while specific constraints may partially shape transmitter co-expression in LNs, a large amount of flexibility on a neuron-by-neuron basis produces heterogeneous network parameters.

As a population, LNs are tonically active (Lei et al., 2011) even without odor-evoked network activity. However, the consequences of LN activation may depend more upon the degree of network activation rather than the identity of any given LN that is activated. Small clear GABAergic vesicles require less calcium influx to trigger fusion relative to dense-core peptidergic vesicles and thus NP release likely only occurs when the network is strongly activated. Additionally, the effects of GABA_B_ receptor activation are far shorter-lasting relative to neuropeptide receptors (Salio et al., 2006), thus NP and GABA co-expression may serve to mediate layered temporal effects on AL neurons. For example, a single modulatory neuron, MCN1, in the stomatogastric ganglion, releases GABA and two neuropeptides which mediate divergent effects with varying time scales, on different neurons in the network (Stein et al., 2007). Furthermore, NP receptors can be excitatory (Horodyski et al., 2011; Lenz et al., 2015; Ormerod et al., 2015) or inhibitory (Yapici et al., 2008; Ignell et al., 2009; Asahina et al., 2014; Ko et al., 2015). All LNs, apart from TKK LNs, co-expressed NPs that activate both inhibitory and excitatory receptors (i.e. MIP and ATR co-expressing LNs), suggesting that the activation of a single LN mediates a complex mix of excitation and inhibition. Sub-populations within each neuronal class likely exhibit differential receptor expression. Thus, future studies will be required to determine if neuropeptide receptor co-expression by individual AL neurons is as heterogeneous as neuropeptide co-expression itself. It may be that the expression of each neuropeptide receptor is regulated by different factors such as current physiological state, as observed for the role of hunger (Ko et al., 2015; Min et al., 2016) or mating state in *Drosophila* (Hussain et al., 2016).

TKK differed in its patterns of co-expression and receptor expression from other neuropeptides. All TKK LNs co-expressed MIP, 84% co-expressed GABA and none co-expressed FMRF or ATR, suggesting that TKK LNs are primarily inhibitory as TKK and MIP receptors are inhibitory in *Drosophila* (Yapici et al., 2008; Ko et al., 2015). Consistent with our data, TKK LNs in the AL of the moth, *Heliothis virescens* also do not co-express FMRF or ATR (Berg et al., 2007). TKK receptor transcripts were not detected in LCC mRNA and thus not in LNs, although TKK LNs could still influence LNs via GABA_B_ and MIP receptor. In *Drosophila melanogaster*, TKK mediates presynaptic gain control upon ORNs (Ignell et al., 2009), and TKKr expression in *M. sexta* ORNs is consistent with this finding. This suggests that TKK LNs may play a distinct role from other LNs in olfactory processing which could include presynaptic gain control. The putative glutamatergic LNs as a population also differed from the GABAergic LNs in that very few non-GABAergic LNs co-express the neuropeptides we examined in this study. Glutamatergic LNs are likely as heterogeneous as GABAergic LNs. Similar to GABAergic LNs, glutamatergic LNs in *Drosophila* are particularly diverse in their morphology (Das et al., 2011) but appear to differ from GABAergic LNs in their synaptic targets by predominantly affecting PNs (Liu and Wilson, 2013), while GABAergic LNs appear to affect all three cell classes, ORNs, LNs and PNs (Wilson and Laurent, 2005; Olsen and Wilson, 2008; Root et al., 2008; Hong and Wilson, 2015).

Our computational simulation suggests that co-expression cannot be explained by random probability alone and adding specific, fixed co-expression relationships, while leaving the rest up to random probability, can replicate the overall co-expression pattern across LNs, perhaps reflecting developmental programs within the AL. Recently the hormone 20-hydroxyecdysone was shown to induce ATR expression in LNs and other neuropeptides in the AL of *M. sexta* (Utz and Schachtner, 2005; Utz et al., 2007). Thus, specific co-expression patterns may reflect developmental cues that guide LN heterogeneity. In cortex, subtypes of local interneurons arise from unique progenitors and their diversity is shaped by additional factors including neural activity and growth factors during development (Huang et al., 1999; Patz et al., 2004; Flames and Marin, 2005). Similarly, GABAergic and glutamatergic LNs in *D. melanogaster*, have been shown to arise from distinct neuroblasts (Das et al., 2008; Das et al., 2011), suggesting that diversity of LNs observed may be, in part, due to distinct origins. However, within subtype diversity may be further shaped by activity dependent mechanisms, like that observed in cortex (Patz et al., 2004). Further study is needed to determine whether similarities in transcriptional control or neural activity may explain the heterogeneity of neuropeptide co-expression patterns.

The high degree of heterogeneity in transmitter co-expression and the overlap in expression of neuropeptide receptors across broad populations of AL neurons, suggest that the dynamic cocktail of peptides present in the AL at any given time likely broadly regulate the modulatory tone of the network. LN-specific co-expression patterns may not functionally matter as long as the network reaches a reliable “set point” which can vary with state-dependent regulation of peptide receptor expression as is the case for hunger (Ko et al., 2015; Min et al., 2016) or mating state (Hussain et al., 2016). Networks have been shown to be tolerant of variable, and diverse parameters with regards to morphology (Otopalik et al., 2017b; Otopalik et al., 2017a), biophysical properties, and synaptic strength (Prinz et al., 2004; Marder and Goaillard, 2006; Marder, 2011), yet still produce robust and consistent output from animal to animal. Similarly, our data suggest that transmitter co-expression from neuron to neuron is flexible, allowing heterogeneous network parameters to emerge and ultimately, expanding the computational capacity of olfactory processing.

## Methods

### Animals

*Manduca sexta* were raised at West Virginia University as previously described (Bell and Joachim, 1976; Daly et al., 2013). Equal numbers of males and females were pooled for all data.

### Immunocytochemistry

Brains were dissected in physiological saline (Christensen and Hildebrand, 1987), fixed in 4% paraformaldehyde overnight at 4°C, embedded in 5% agarose to be sectioned at 100µm using a Leica VT 1000S vibratome. Sections were washed in PBS with 1% triton X-100 (PBST), blocked in PBST and 2% IgG free BSA (Jackson Immunoresearch; Cat#001-000-161) and then incubated in blocking solution with 5mM sodium azide and primary antibodies. For all rabbit-neuropeptide/mouse-GABA protocols, sectioned tissue was incubated for 48 hours at dilutions of 1:3000 and 1:500 respectively. The mouse-BRP antibody was used at a dilution of 1:50 and incubated at 4°C for 172 hours. Sections were then briefly washed with PBS, PBST, cleared with ascending glycerol washes and then mounted on slides with Vectashield (Vector Laboratories; Cat#H-1000). All neuropeptide antibodies used in this study were raised in rabbit. For protocols in which we labeled with two antisera raised in rabbit, we used APEX Antibody Labeling Kits 488, 555, 647 (Invitrogen;Cat #s A10468, A10470, A10475, respectively) to directly attach a fluorophore with excitation/emission spectra at different wavelengths to each primary to avoid cross-labeling from a secondary antibody (Bradley et al., 2016). Using the resin tip from the APEX kit, a small amount of the antibody (10-20µg) was pushed through the resin using an elution syringe and the reactive dye was prepared using DMSO and a labeling buffer (Solutions provided in APEX kit). The reactive dye was eluted through the tip onto the antibody remaining in the resin to covalently bond the fluorescent label to the IgG antibodies. The tip was incubated overnight 4◦C or at room temperature for 2 hours and the labeled product was eluted through the tip. Resulting labeled antibody volume of 50uL in a total volume of 2400ul was used to label 6 brains at equal dilution of 400ul per well and incubated for 72 hours in 3% triton X-100 with PBSAT. Sections were then washed and mounted as above.

### Antibody Characterization

Specificity controls (including pre-adsorption controls and western blots analysis) for the AST, ATR, TKK, MIP and BRP antibodies in *M. sexta* brain tissue are described in (Lizbinski et al., 2016). GABA pre-adsorption controls in *M. sexta* AL tissue for the mouse GABA antiserum are described in (Bradley et al., 2016).

BRP-The BRP antiserum (nc-82: Developmental Studies Hybridoma Bank) was raised in mouse against the *Drosophila melanogaster* protein “Bruchpilot” which is a homologue of the presynaptic active zone protein ELKS/CAST and is required for functional synapses in the nervous system of *Drosophila* (Wagh et al., 2006). We used the BRP antibody to delineate glomerular boundaries within the AL of *M. sexta*. (RRID: AB_2314866)

GABA- The GABA antibody (Sigma Aldrich, cat # A2052) was raised in rabbit against GABA coupled to BSA with paraformaldehyde.

MIP- Antiserum raised in rabbit against MIP conjugated to thyroglobulin was produced by M. Eckert, Jena Germany and provided by C. Wegener, Marburg Germany (Predel et al., 2001). (RRID: AB_2314803).

ATR- Antiserum raised in rabbit against *M. sexta* allatotropin (Mas-AT, referred to in this paper as ATR) was kindly provided by Dr. J. Veenstra, University of Bordeaux, Talence, France; (Veenstra and Hagedorn, 1995). (RRID: AB_2313973)

AST-A- Antiserum was raised (Reichwald et al., 1994) in rabbit against octadecapeptideallatostatin (Pratt et al., 1991), ASB2, (AYSYVSEYKALPVYNFGL-NH2) of *Diploptera punctata* and kindly provided by Dr. J. Veenstra, University of Bordeaux, Talence, France. It recognizes AKSYNFGLamide, a form of AST and other AST-like peptides.

TKK- Antiserum raised in rabbit against locust tachykinin II with bovine thyroglobulin with glutaraldehyde was kindly provided to us Dr. J. Veenstra, University of Bordeaux, Talence, France. (RRID: AB_2341129)

FMRF- FMRFamide antiserum was raised against synthetic RF-amide coupled to bovine thyroglobulin with glutaraldehyde and provided by Dr. Eve Marder (Marder et al., 1987). Pre-adsorption controls of the antiserum against synthetic FMRFamide eliminated labeling in larval Manduca nervous tissue (Witten and Truman, 1996).

### Confocal Microscopy

Image stacks were scanned using an Olympus Fluoview FV1000 confocal microscope with argon and green and red HeNe lasers. Fluoview was also used to set brightness levels and Corel Draw X4 was used to organize figures.

### Cell counts and co-expression

Images of immunostained brains were exported as .tiff stacks in Fluoview software. Stacks were then imported into VAA3D software to determine individual cell counts and co-localized cell counts. The number of local interneurons in the LCC that express each transmitter were counted in VAA3D (n=6 brains per label combination). We used cell body size, and location within the LCC to distinguish between LNs and PNs (Homberg et al., 1988). The average and standard deviation of number of cells per AL that expressed a given transmitter were calculated for each combination. Wilcoxon rank sum tests were performed using Graph Pad Prism v.6.01 (Graphpad Software Incorporated) to determine if there was any significant difference between the left and the right AL for each brain. Co-expression ratios were determined by dividing the number of cells expressing both an individual neuropeptide and GABA by the total number of cells expressing just the neuropeptide and calculated in Excel. Neuropeptide co-expression ratios were determined in the same manner for every possible pairwise combination using data from direct labeling experiments. FMRF/MIP co-expression ratios were not calculated as the direct tips labeled significantly less MIP neurons than all other runs and therefore the ratios would not have reflected accurate co-expression. Thus, FMRF/MIP co-expression was not used in subsequent models or computational analysis as neither a constraint nor a relationship to replicate. All other neuropeptide/neuropeptide co-expression experiments labeled an accurate # of cell bodies when compared to cell counts from GABA/neuropeptide runs using indirect immunocytochemistry. Cell count totals and standard deviations from direct labeling were used in all model iterations, as co-expression ratios were calculated using that data.

### Putative neuropeptide receptor sequence BLAST

We used receptor sequences from closely related invertebrate species to identify putative sequence homologs on *M. sexta* scaffolds. Protein sequences from *Drosophila melanogaster* and other closely related species were identified by annotation (see Table 2) and queried against the *M. sexta* genome using tblastn (National Agricultural Library, i5k initiative https://i5k.nal.usda.gov/Manduca_sexta). Top matches to each receptor sequence in *M. sexta* were subsequently queried against the NCBI nr database to confirm their putative annotation as *M. sexta* receptor homologs. These sequences were used for primer design for RT-qPCR analysis of putative neuropeptide receptor expression in the antennae, medial and lateral cell clusters, and brain. Sequences that were previously identified in *M. sexta* for ATRr and RpS3 (Jiang et al., 1996; Horodyski et al., 2011) were downloaded as FASTA files from NCBI (http://www.ncbi.nlm.nih.gov/gene/?term=) and used to design RT-qPCR primers. Open reading frames were established using ORF Finder at http://www.ncbi.nlm.nih.gov/projects/gorf/. A recent study partially annotated the *M. sexta* genome (Kanost et al., 2016). We used the *M. sexta* raw sequence, and assembled genome sequence at NCBI Assembly ID GCA_000262585 from Kanost (http://www.ncbi.nlm.nih.gov/assembly/GCA_000262585.1) (Kanost et al., 2016) and identified the sequence IDs for each of the transcripts in question (Table 1). None of the putative receptor sequences are currently annotated in NCBI Assembly ID GCA_000262585.

### Primer design

Open reading frame nucleotide sequences for each receptor, as established above, were used as the basis for primer design for RT-qPCR. Primers were designed using http://www.bioinformatics.nl/cgi-bin/primer3plus/primer3plus.cgi and checked for optimal conditions using OligoAnalyzer 3.1 (https://www.idtdna.com/calc/analyzer). Primers and amplicons were then run through a BLAST of the *M. sexta* genome to determine if they matched to the specified sequence and to rule out potential priming mismatches with other parts of the genome. Table 1 lists primer sequences and annealing temperatures. All primers used for RT-qPCR amplified a 90-125bp stretch of sequence.

### Primer design

Open reading frame nucleotide sequences for each receptor, as established above, were used as the basis for primer design for RT-qPCR. Primers were designed using http://www.bioinformatics.nl/cgi-bin/primer3plus/primer3plus.cgi and checked for optimal conditions using OligoAnalyzer 3.1 (https://www.idtdna.com/calc/analyzer). Primers and amplicons were then run through a BLAST of the *M. sexta* genome to determine if they matched to the specified sequence and to rule out potential priming mismatches with other parts of the genome. Table 1 lists primer sequences and annealing temperatures. All primers used for RT-qPCR amplified a 90-125bp stretch of sequence.

### Real-time(RT) qPCR

Antennae, medial cell clusters, lateral cell clusters and brains were collected from 2-6 day old *M. sexta* and RNA was extracted using a TRIzol reagent (Molec. Research center Cat# # TR 118). Equal numbers of pooled males and females were used for each biological tissue sample for a total of 3 biological samples for each tissue type (n=3 antennae; n=40 medial cell cluster from 20 brains; n = 40 medial cell cluster from 20 brains; and n= 2 brains). We used the 40s ribosomal protein s3 (RpS3) as our reference gene. RpS3 expression values were consistent across biological replicates. RNA was treated with TURBO DNA-free™ Kit (ThermoFisher Scientific Cat# AM1907) to prevent genomic DNA contamination and cDNA was synthesized using the SuperScript^®^ IV First-Strand Synthesis System (ThermoFisher Scientific Cat#18091050). We performed RT qPCR with the BioRad CFX Connect Real-Time System (Cat #1855201) to determine the relative expression of putative neuropeptide receptors across our tissue samples. Individual samples were prepared by combining prepared cDNA sample, [100um] forward and reverse primers, SsoFast EvaGreen Supermix (BioRad Cat # 1725200) and nuclease free diH_2_0 to a volume of 10ul. RT^−^ samples, no template controls (NTCs) and positive controls with *M. sexta* genomic DNA from the brain were run for every plate. RT^−^ and NTC had no amplification for all receptors and sample types run at 58.3°C (See Table 3). Optimal annealing temperatures were determined through a gradient test on genomic DNA to ensure qPCR on cDNA was performed at optimal temperature. All primer sets, including the reference gene, RpS3, were run using the following protocol (95°C 2 minutes, (95°C 5 seconds-> 58.3°C 30 seconds) x 39 cycles, 65.0°C 5 seconds stepped up to 95°C) except for ATRr primers, which were annealed at a temperature of 52°C. All samples for for RpS3 were run again at 52°C to ensure accurate calculation of relative expression values for ATRr. Cq values for ANTa (antennae sample a), Mb (Medial cell cluster sample b) and NTC sample for the RpS3 run at 52°C were high (see Table 3). However, amplification curves revealed that there were no sharp amplification peaks and thus, high Cq values were due to noise not contamination. High Cq values with non-descript peaks for RpS3 NTCs run at 52°C were considered 0 for ANTa, Mb and NTC when calculating relative expression.

**Table 3:** Cq values for all receptors and RpS3 from RT-qPCR

|  |  | ANTa | ANTb | ANTc | Ba | Bb | Bc | Ma | Mb | Mc | La | Lb | Lc | genomic |
| --- | --- | --- | --- | --- | --- | --- | --- | --- | --- | --- | --- | --- | --- | --- |
| TKKr | RT | 0 | 34.1 | 34.5 | 31.36 | 35.35 | 29.63 | 38.51 | 32.18 | 34.03 | 39 | 0 | 37.59 | 24.09 |
|  | RT- | 0 | 0 | 0 | 0 | 0 | 0 | 0 | 0 | 0 | 0 | 0 | 0 | 0 (NTC) |
| ATRr | RT | 35.72 | 33.65 | 33.41 | 32.28 | 37.32 | 31.26 | 39.8 | 32.42 | 33.55 | 35.77 | 37.6 | 36.29 | 30.24 |
|  | RT- | 0 | 0 | 0 | 0 | 0 | 0 | 0 | 0 | 0 | 0 | 0 | 0 | 39.48 (NTC) |
| FMRFr | RT | 32.43 | 29.04 | 29.1 | 26.56 | 31.14 | 25.95 | 33.01 | 27.53 | 28.03 | 31.57 | 32.22 | 30.89 | 24.85 |
|  | RT- | 0 | 0 | 0 | 0 | 0 | 0 | 0 | 0 | 0 | 0 | 0 | 0 | 0 (NTC) |
| MIPr | RT | 33.91 | 31.03 | 30.69 | 29.55 | 33.31 | 27.94 | 34.94 | 28.92 | 30.15 | 34.97 | 35.73 | 33.12 | 23.86 |
|  | RT- | 0 | 0 | 0 | 0 | 0 | 0 | 0 | 0 | 0 | 0 | 0 | 0 | 0 (NTC) |
| ASTr | RT | 37.28 | 32.23 | 31.23 | 29.26 | 23.63 | 28.17 | 37.39 | 31.19 | 32.39 | 35.43 | 37.35 | 35.14 | 25.61 |
|  | RT- | 0 | 0 | 0 | 0 | 0 | 0 | 0 | 0 | 0 | 0 | 0 | 0 | 0 (NTC) |
| GABA<sub>B</sub> | RT | 32.21 | 29.09 | 28.62 | 27.91 | 31.17 | 25.97 | 33.48 | 27.26 | 28.51 | 33.26 | 35.51 | 31.97 | 23.72 |
|  | RT- | 0 | 0 | 0 | 0 | 0 | 0 | 0 | 0 | 0 | 0 | 0 | 0 | 0 (NTC) |
| RpS3 | RT | 25.61 | 22.49 | 22.16 | 22.91 | 26.65 | 21.41 | 28.89 | 23.16 | 24.04 | 28.93 | 28.6 | 26.04 | 24.18 |
|  | RT- | 0 | 0 | 0 | 0 | 0 | 0 | 0 | 0 | 0 | 0 | 0 | 0 | 0 (NTC) |
| RpS3<br>(at 52°C) | RT | 24.54 | 21.85 | 21.87 | 23.18 | 27.04 | 21.31 | 28 | 23.21 | 24.18 | 26.68 | 28.03 | 25.23 | 18.29 |
|  | RT- | 4.41 | 0 | 0 | 39 | 37.27 | 0 | 37.2 | 5.13 | 28.94 | 36.16 | 38.98 | 0 | 5.23 (NTC) |

### qPCR relative expression analysis

Raw qPCR data can be found in Table 3. Delta Ct (Ct_receptor_-Ct_reference gene_) values were calculated for each receptor using RpS3 Ct values as the reference gene, and averaged across all biological replicates for brain, LCC, MCC and antennae tissue samples. Relative expression levels (2^−ΔCT^) were calculated for all receptors. Ct values less than or equal to 37 were considered non-detectable. All graphical representations for receptor qPCR were performed in GraphPad prism (v. 5.01).

### Computational analysis of transmitter co-expression

We wrote a MATLAB (Naticks MA) program to determine if our observed transmitter co-expression data could be replicated by random probability. Given the known total number of LNs in the LCC, and the total number of LNs expressing each neuropeptide from our cell counts, the model calculates the random probability of a given neuron co-expressing two transmitters. The program is given the average number of cells expressing a given neurotransmitter and then randomly assigns them to one of the cells in the cluster. The model can thus calculate the probability of pairwise co-expression (i.e. 100% of TKK cells express MIP) in the lateral cell cluster based on chance. Specifically, using our biological data as the backbone of the model, we designed a matrix with 6 columns, 1 per transmitter type (TKK, FMRF, ATR, MIP, AST, GABA), with a row length of 360 long (the total number of LNs in the lateral cell cluster). Within each column, the model randomly distributes the number of cells that express a given transmitter to a row between 1 and 360 (Supp. Fig 1). For example, if we know that 12 cells within the LCC are TKKergic, the model randomly picks 12 numbers between 1 and 360 in the TKK column, and marks that cell as TKK positive. The number of cells expressing a certain neurotransmitter is chosen probabilistically based on the observed average and standard deviation. With each iteration of the model, the cells that are assigned as ‘transmitter positive’ within each column are randomized. The model does this with all respective cell count totals for each transmitter column and then calculates the percentage each transmitter will be expressed with another transmitter by pure random chance (across all pairwise comparisons) 10,000 times. Standard deviation and percentages of co-expression were recorded from our model’s output. We then compared our known co-expression percentages to the model’s output to determine if the biological data reflected a pattern above random chance.

The model described above has no initial assumption about the likelihood of co-expression and only the overall number of cells expressing each of the transmitter is determined initially. We used a similar program to determine if certain co-expression relationships were predictive of other co-expression relationships within the cell cluster. To do this, we built certain co-expression relationship explicitly as initial assumptions. For example, if we know that on average 97.2% of those ATRergic cells are also GABAergic, the program explicitly chose 97.2% of the cells that where assigned to be ATR positive to also be GABA positive. This co-expression relationship is thus no longer determined at random like the first version of the script, but rather is an initial assumption – a rule. We can then determine if this rule alone shifts the co-expression of other transmitter types closer to the known biological co-expression percentages. We applied these rules one by one, for every pairwise comparison of co-expression and statistically compared the output of the random model to the output of the rule model as well as our known biological data. This allowed us to determine if specific co-expression relationships replicate other co-expression relationships.

MatLab script available at www.dacksneuroscience.com

### Data analysis

Standard deviation indices (SDIs) were calculated to determine how well each model predicted the probability of transmitter co-expression. Similar to a Z-score, this measure determines how close the model’s predicted probability is to the known biological probability of co-expression.

SDI = (Mean_model_ – Mean_biological_)/stdev_greatest_

Where Mean_model_ = Mean probability of co-expression of any two given neurotransmitters from the model e.g. mean % TKK co-localized with MIP

Mean_biological_ = Mean probability of co-expression of any two given neurotransmitters from the co-expression cell counts

stdev_greatest_ = the greatest standard deviation from either the model or biological data for a given co-expression relationship.

Weighted SDI’s were calculated to take the number of neurons which expressed each transmitter into account.

Weighted SDI = ∑ ((Mean_model_ – Mean_biological_)/stdev_greatest_)* (# of co-localized cells predicted/total # of LCC cells)

For example, there are only 12 TKK neurons in a total of 360 LNs, but 142 ATR neurons. Therefore, predicting the # of ATR neurons vs. TKK neurons should carry more weight when determining the accuracy of each model. Weighted SDIs for each co-expression relationship (i.e. weighted SDI for the TKK/MIP co-expression) were summed across relationships for an overall measure of the accuracy with which each model iteration replicated observed co-expression patterns.

SDI values can be interpreted by the following scale:

0: perfect consensus between model and experimental data

1: model results are within one standard deviation of experimental data and thus replicate the data reasonably well;

2: model results are within two standard deviation of experimental data and thus do not replicate the data accurately;

To determine the % improvement of each model at replicating known biological co-expression (Fig 3G), all weighted SDI’s were normalized with respect to the weighted SDI of the random model using the formula:

% improvement from random = (1-(weighted SDI_x_/weighted SDI_random_))*100

All statistics were performed in GraphPad prism (v. 5.01).

## Acknowledgements

We would like to thank Dr. Tim Driscoll and Dr. Tori Verhoeve for their advice and help with RT-qPCR experimental design and setup, Andrew Steele and Lillian Bailey for their help with cell counts and Aditya Kesari for assistance with preliminary data as well as the other members of the Dacks lab for their support. We would also like to thank Kate Allen, Tyler Sizemore, Phillip Chapman, Dr. Kevin Daly, Dr. Sadie Bergeron and Dr. Quentin Gaudry for helpful comments on the manuscript. The TKK, AST, and ATR antibodies were provided by Dr. Jan Veenstra, the MIP antibody was provided by Dr. Christian Wegener and developed by Dr. Manfred Eckert and the FMRF antibody by Dr. Eve Marder. This work was supported by start-up funds from West Virginia University, an R03 DC013997-01 from the National Institutes of Health, and USAFOSR FA9550-17-1-0117 to AMD.

## Author Contributions

All authors had full access to all the data in the study and take responsibility for the integrity of the data and the accuracy of the data analysis. Study concept and design: Kristyn M. Lizbinski (KML), Gary F. Marsat (GFM), Andrew M. Dacks (AMD). Acquisition of data: KML Computational model: KML, GFM. Analysis and interpretation of data: KML, GFM, AMD. Drafting of the manuscript: KML, GFM, AMD. Critical revision of the manuscript for important intellectual content: KML, GFM, AMD. Obtained funding: AMD Administrative, technical, and material support: AMD. Study supervision: AMD

---

**Supplemental Figure 1:**
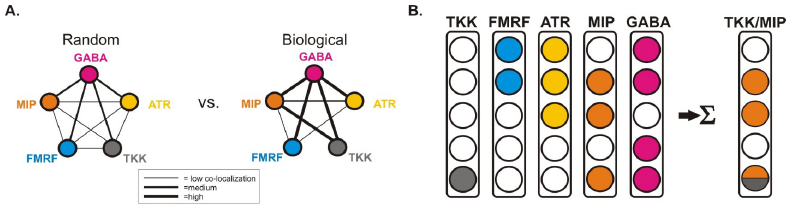
Calculating the probability of LN co-expression patterns using computational modeling. **A:** Schematic representation of random vs. biological co-expression. Each circle represents a population of LNs that express a given transmitter. Given the expression values in Fig. 1F, the random likelihood that a neuron will express two transmitters is calculated and represented by line thickness. **B:** Schematic representation of modeling procedure. Each column represents a transmitter, the number of rows corresponds to the total # of neurons in the population (reduced in this illustration to 5 total cells for the sake of simplicity). The model then sums across each row in a pairwise fashion to determine the co-expression of a given transmitter pair. The probability that a given neuron will co-express a given pair of transmitters is output as a percentage.

